# A Subtractive Proteomics Approach for the Identification of Immunodominant *Acinetobacter baumannii* Vaccine Candidate Proteins

**DOI:** 10.1101/2022.06.26.497689

**Authors:** Mustafa Burak Acar, Şerife Ayaz-Güner, Hüseyin Güner, Gökçen Dinç, Ayşegül Ulu Kılıç, Mehmet Doğanay, Servet Özcan

## Abstract

*Acinetobacter baumannii* is one of the most dangerous multidrug-resistant pathogens worldwide. Currently, 50-70% of clinical isolates of *A. baumannii* are extensively drug-resistant (XDR) and available antibiotic options against *A. baumannii* infections are limited. There are still needs to discover specific *de facto* bacterial antigenic proteins that could be effective vaccine candidates in human infection. With the growth of research in recent years, several candidate molecules have been identified for vaccine development. So far, there is no public health authorities approved vaccine against *A. baumannii*. The purpose of this study was to identify immunodominant vaccine candidate proteins that can be immunoprecipitated specifically with patients’ IgGs. Relaying on hypothesis that IgGs of infected person have capacity to capture immunodominant bacterial proteins. Herein, outer membrane and secreted proteins of sensitive and drug resistant *A. baumannii* were captured by using IgGs obtained from patient and healthy control sera and were identified by LC-MS/MS analysis. By using subtractive proteomic approach, we determined 34 unique proteins which were captured only in drug-resistant *A. baumannii* strain via patient sera. After extensive evaluation of predicted epitope regions, solubility, membrane transverse characteristics, and structural properties, we selected several notable vaccine candidates. We identified vaccine candidate proteins that triggered *de facto* response of human immune system against the antibiotic-resistant *A. baumannii*.

## Introduction

*Acinetobacter* spp. are gram-negative, aerobic, non-fermentative, stabile, and bacillus-shaped bacteria, reside in sand and water. Approximately 25% of healthy people can contain these bacteria in their armpit, groin, and even in the oral cavity and respiratory track[1,2]. This species forms the pinky colonies on Mac Conkey agar, can be shown as bacillus, coccobacillus and they can survive even in dry conditions[3].

Following the surgical operations, insertion of an intravascular, ventricular catheter or endotracheal tube can bring risk of nosocomial infection occurrence. *Acinetobacter baumannii* can surround the wound site or damaged mucosal areas. Being an opportunistic bacterium, *A. baumannii* mainly infects critically-ill patients hosted in intensive care units (ICU)[4]. Despite the treatment with combined antibiotics, emerging multi drug resistance makes the treatment ineffective and infection caused by these bacteria can result in death[5].

*A. baumannii* is one of the most dangerous multi-drug resistant pathogens all around the globe. Currently, 50-70% of *A. baumannii* clinical isolates have extensive drug resistance (XDR) and the frequency of the infections caused by these bacteria are increasing[6,7]. By the development of resistance against colistin and tigecycline stands for *A. baumannii* may become a pan drug resistant (PDR) bacterium which causes the infection that incurable with FDA approved antibiotics[8,9]. Running out of antibiotic options that is available against *A. baumannii* reveals the necessities of vaccination and development of alternative treatment approaches against this bacterium. With the increasing number of studies in recent years, several candidate molecules have been revealed for vaccine development. The antigens use in these studies are including outer-membrane proteins, plays an important role in virulence and biofilm genesis of this bacterium, such as OmpA, Omp34, OprC Phospolipase C and D[5,10–14]. Especially -omics and *in silico* techniques with the potential of generating and analyzing the high-throughput data identified candidate vaccine molecules. Unfortunately, as against other opportunistic bacterial infections, there isn’t an available vaccine approved by health authorities for *A. baumannii* [15]. An ongoing need for effective vaccine candidate still in search to specifically target *de facto* molecules in human infection.

This study aimed to identify those potential vaccine candidate proteins which can be specifically immunoprecipitated with the IgGs that present only in patients suffering from drug resistant *A. baumannii* infections. Therefore, outer membrane and secreted proteins of sensitive and drug resistant bacteria were captured by using immunoglobulin containing patient and control sera and were identified by LC-MS/MS analysis. To determine patient specific ones, we excluded bacterial antigens that immunoprecipitated by control sera. Our approach allowed us to identify 34 unique bacterial proteins that triggered immune response in infected individuals, but not in the control and healthy individuals. By performing bioinformatic evaluations, epitope prediction, solubility, transmembrane properties and structural specifications, selected 9 proteins were further evaluated for possible recombinant applications. Hence, we identified several candidate molecules which could have the potential protective role against *A. baumannii* infections.

## Results

In this study, we have used blood sera of 29 *A. baumannii* infected intensive care unit patients. As control 13 individuals who were treated in the same intensive care unit, with having no *A. baumannii* infection or any other bacterial infection. Additionally, three healthy individuals were considered as the external control group to eliminate the conserved antigens originated from normal flora members.

Bacterial surface proteins and secretome proteins were isolated from multiple drug resistant (BAA-1710) and non-resistant (ATCC 17978) *A. baumannii* strains and used to immune interaction with human sera. In order to capture mutual/conserved antigens, we used standard ATCC bacterial strains instead of unmapped local clinical isolates. To minimize biological variation between individuals, the sera of the same blood type were combined before immunoprecipitation. The workflow of our experimental strategy was depicted at Figure 1.

**Figure 1.**
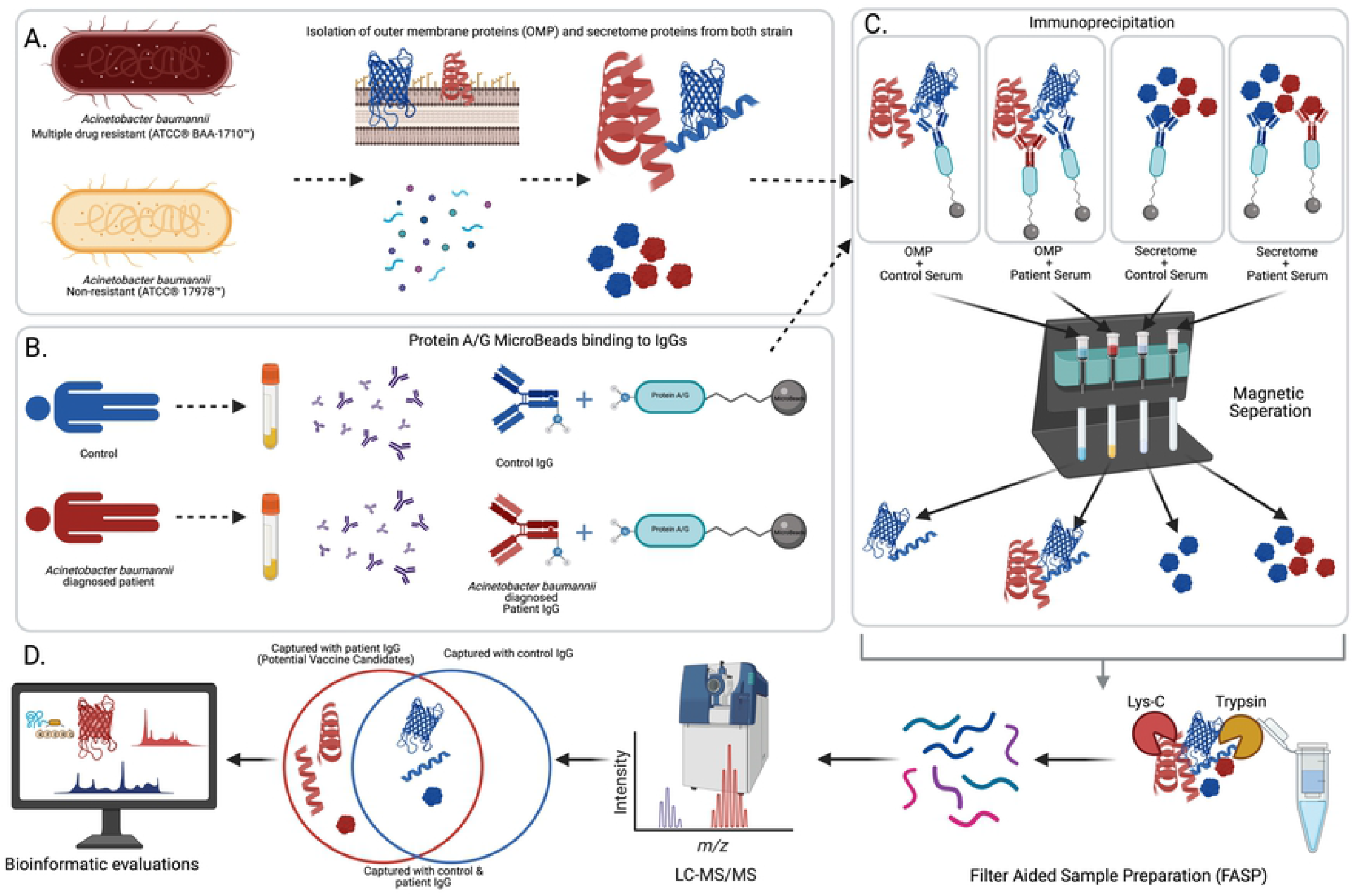
The workflow of our experimental strategy. **A**. Isolation of outer membrane and secretome proteins of *Acinetobacter baumannii* resistant (BAA-1710) and sensitive strain (17978). **B**. IgGs from human sera were linked to Protein A/G Microbeads. **C**. Protein A/G bound IgG were utilized to bait immunodominant *A. baumannii* proteins from each group. **D**. Sample preparations, LC-MS/MS analysis and bioinformatic evaluations of identified antigenic proteins. Created with BioRender (https://biorender.com/).

### Identified subcellular immunogenic proteins of A. baumannii

LC-MS/MS analysis that followed by immunoprecipitation with patient sera, revealed 49 outer membrane proteins of resistant strain (BAA1710). When immunoprecipitation was performed with control sera, 41 immunogenic proteins were identified for the same strain. Thirty-four of these were commonly identified from control and patient sera, whereas 15 antigens were specifically captured only with patient sera and accession numbers presented in **Figure 2A**. To reason these 15 proteins specific to resistant strain we have mapped sensitive strain (17978), we identified 23 outer membrane protein by immunoprecipitation with control sera, 13 proteins with patient sera. Comparison of these two groups showed, only two of these proteins were specifically precipitated with patient sera **(Figure 2B)**, but none of these proteins were shared with resistant strains finding; therefore 15 immunogenic outer membrane proteins were classified only to the resistant strain.

**Figure 2.**
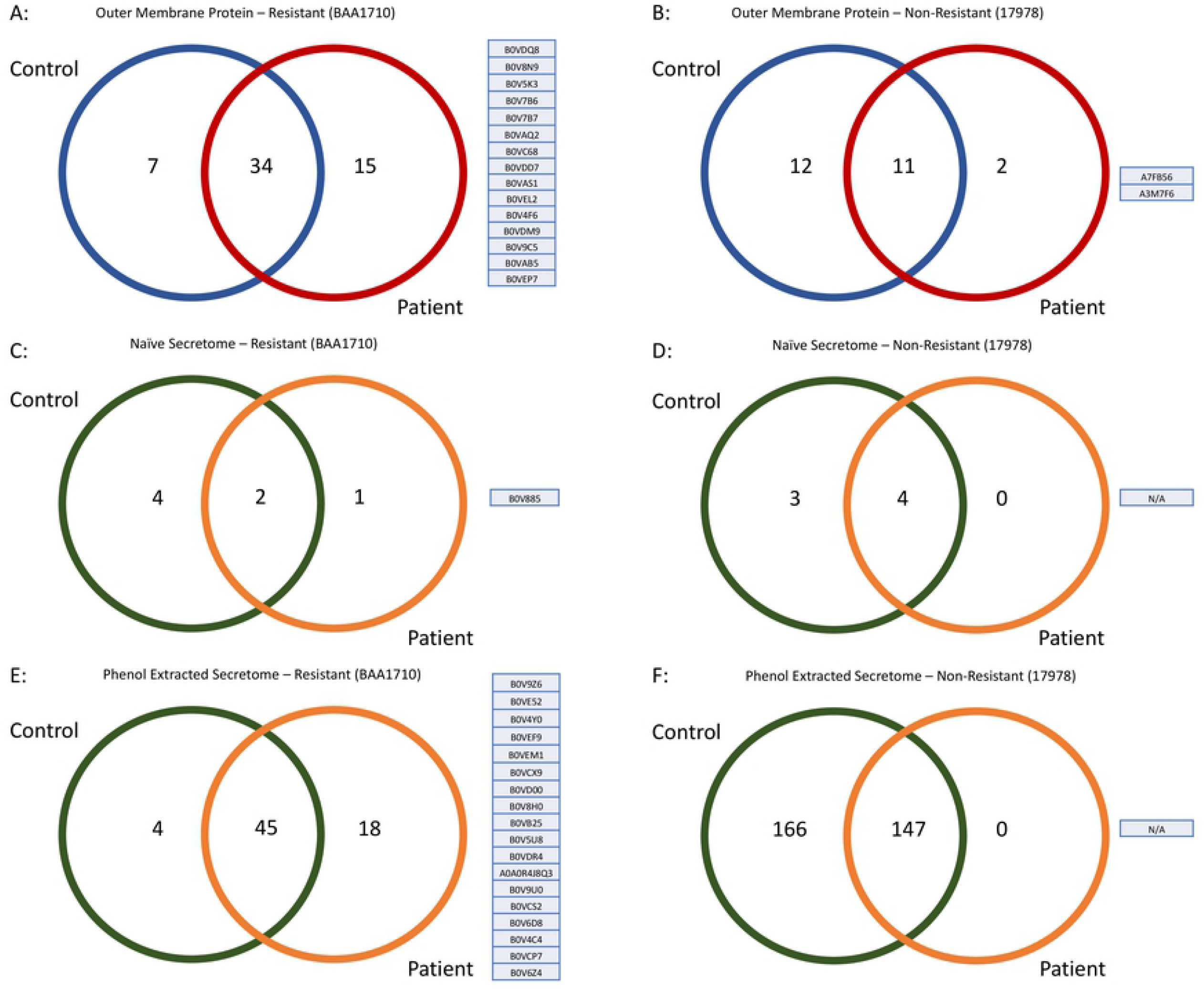
Venn diagram and accessions numbers of identified proteins from different groups. **A**. 15 outer membrane proteins of resistant strain that captured via patient sera and their accession numbers. **A**. 2 outer membrane proteins of sensitive strain that captured via patient sera and their accession numbers. **C**. One secretome proteins of resistant strain that captured via patient sera without any enrichment protocol and its accession number. **D**. No antigenic secretome protein of sensitive strain were captured with patient sera in naïve form. **E**. 18 secretome proteins of resistant strain that captured via patient sera after enrichment with phenol extraction and their accession numbers. **F**. No antigenic secretome protein of sensitive strain were captured with patient sera after phenol extraction.

We immunoprecipitated bacterial proteins directly from conditional medium, without any protein enrichment protocols (i.e., phenol, TCA-Acetone precipitation, etc.), from both strains with control and patient sera. Accomplished LC-MS/MS analysis revealed that only one immunogenic naive secretome protein of resistant strain was captured and identified with patient sera **(Figure 2C)**, while there wasn’t identified protein from naïve secretome of sensitive strain **(Figure 2D)**. The only protein that was captured B0V885. This made us think that it should be highly abundant in conditioned medium, therefore we were capable to capture it without any pre-processes. Since very low number of proteins were identified from naïve secretome, we concentrated secretome proteins with the phenol-chloroform precipitation. By doing immunoprecipitation with patient sera to phenol precipitated secretome proteins, we identified 18 proteins specific to resistant strain **(Figure 2E)** while there were not any identified proteins of sensitive strain **(Figure 2F)**.

A total of 34 antigenic proteins were identified from the outer membrane and secretome (naïve and phenol precipitated) via immunoprecipitation with patient sera, specifically belonging to the resistant strain **(Table 1)**. Majority of identified proteins were bacterial membrane and periplasm. When evaluated their molecular function, some molecules functions as membrane transporter, symporter, ion binding, antibiotic binding, outer membrane assembly and protein secretion. Also, some molecules have enzymatic activity like hydrolase, oxidoreductase and cis-trans isomerase.

### B-Cell epitope prediction

We used a sequence-based epitope prediction tool, IEDB in the prediction of identified proteins B-Cell epitopes. Outputs of this tool showed us that B0V885 and B0VAB5 have highly scored epitope regions. Also, the rest of the immunogenic proteins had a nonnegligible epitopic region score **(Supplementary File 1)**.

### Structural and solubility predictions of identified proteins

Predictions by BOCTOPUS revealed that 9 of identified proteins have transmembrane beta-barrel structure topology. These molecules were B0V4F6, B0V5K3, B0V7B6, B0V7B7, B0V9C5, B0V9U0, B0VCP7, B0VE52, B0VEM1. These phenomena were evaluated with PHYRE2 and it showed us that B0V7B6, B0VCX9, B0VEM1 and B0V9U0 have also beta-barrel structure. By viewing two different logic we were able to identify total of 13 out of 33 proteins demonstrating beta-barrel structure, which presumably insoluble. One of the identified proteins B0VEF9 had a quite long amino acids length (8200 aa), we were not able to get any prediction within the limitation of both tools. We also evaluated the transmembrane properties of identified proteins, TMHMM tool provided us that, B0VDQ8, B0V8N9, B0VDD7, B0VDM9, B0V9C5, B0VEP7 had ≥2 transmembrane regions. Additionally, solubility scores of the identified proteins assessed with PROSO II and proteins, which had a score above 0.6 were accepted as soluble. Of the 34 proteins, only 8 proteins were above the threshold of solubility with the default parameters (B0V7B7, B0VC68, B0V4F6, B0VAB5, B0V9Z6, B0VE52, B0VD00, B0V8H0, A0A0R4J8Q3).

### Possible candidates for recombinant vaccine production

Based on the structural analysis and solubility scores, we mainly focused on eight predicted soluble proteins as a candidate. Additionally, we evaluated the rest of our list of proteins on immunogenicity and epitope predictions. Even though B0V885 was detected as insoluble by PROSO II, when we scrutinized IEDB outputs, it was the highest epitope conveying protein. It is of note, B0V885 was uniquely captured from naïve secretome, without any enrichment, in the native conformational state. Furthermore, when the reference genomes of resistant and sensitive strains were aligned, a noticeable peptide region (between the 735^th^ and 752^nd^ amino acids of B0V885 protein) was present in the resistant strain but not insensitive strain **(Figure 3)**. Above all those reasons, B0V885 had attracted our attention as a druggable protein. Therefore, our recombinant vaccine candidate list rolled up to nine.

**Figure 3.**
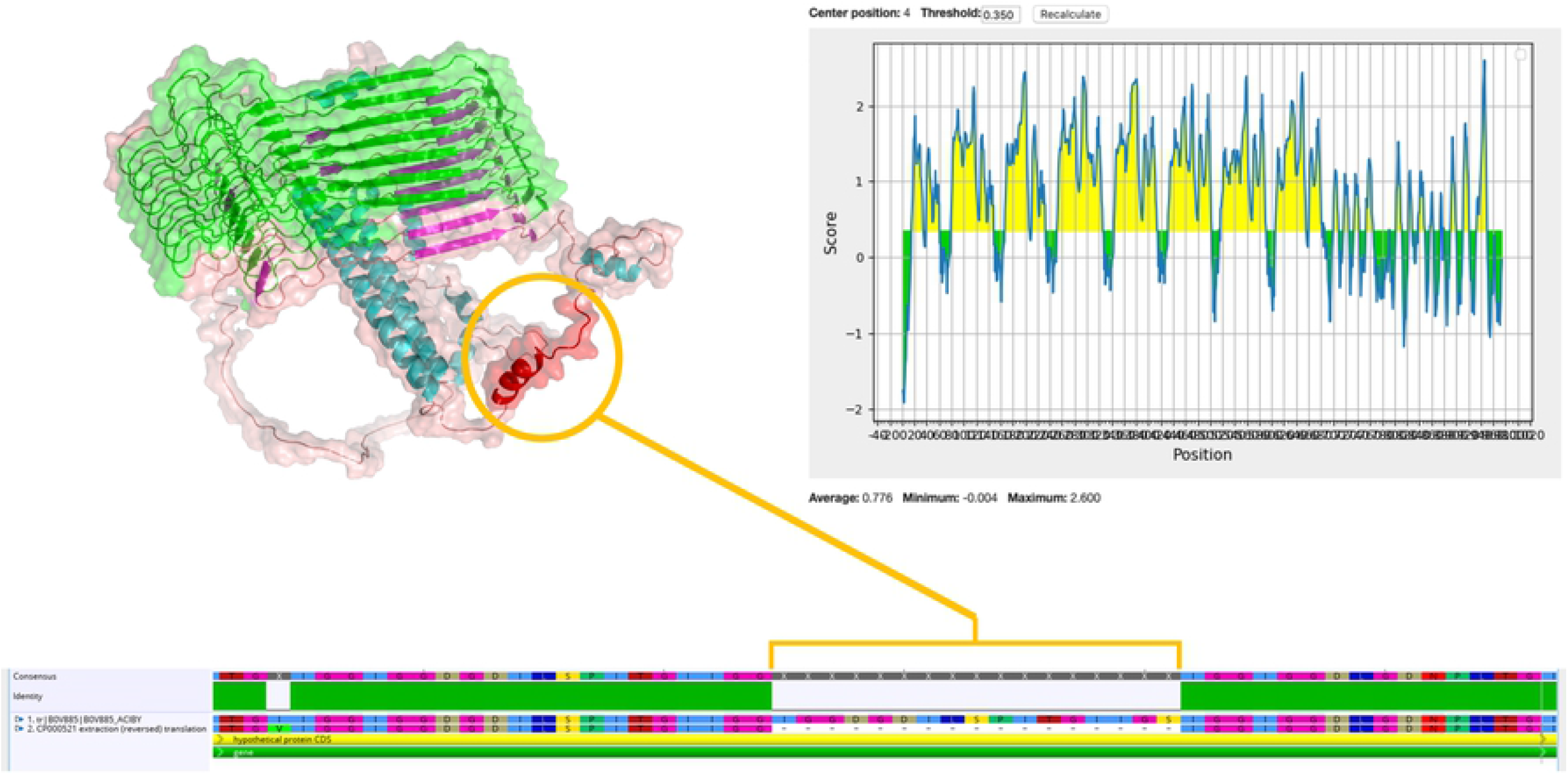
Predicted epitope regions and 3D structure of B0V885 protein. Circled red helix was a 17 amino acid length region that present in the resistant strain whereas it was not present in the sensitive.

### Epitope alignment to protein structure

The expectation was an epitope was valid if it was located on accessible region protein. Therefore, we wonder if the nine selected were fitting into this criterion. State-of-the-art visualization of the candidate proteins we employed AlphaFold2. We successfully calculated three-dimensional models of our nine proteins **(Figure 4)**. Thanks to AlphaFold2 for enabling us to model such large proteins which was not successful with other prediction tools. On the three-dimensional models of our proteins, selected epitopic regions (e.g., length of the epitopic region and possible O-linked Glycosylation) were highlighted. Also, Ramachandran plots were extrapolated to verify AlphaFold2 generated three-dimensional models, if it was in favorable regions of the plots.

**Figure 4.**
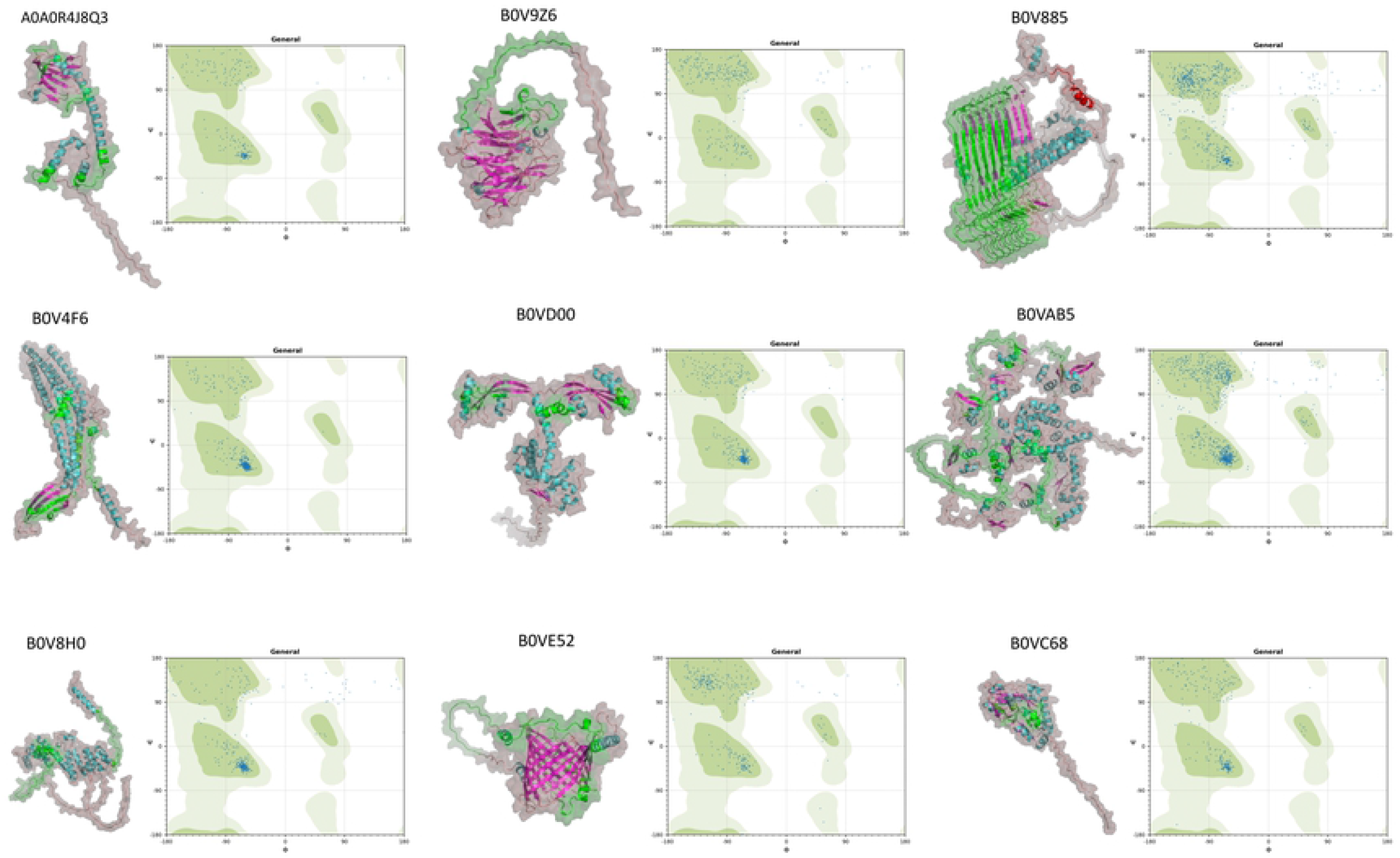
Tertiary structure prediction of nine selected proteins (A0A0R4J8QA3, B0V9Z6, B0V885, B0V4F6, B0VD00, B0VAB5, B0V8H0, B0VE52, B0VC68) by AlphaFold v2.1.0. IEDB predicted B-Cell epitope regions were overlapped and highlighted with green labelling. Ramachandran Plots of these proteins were also depicted for the presentation of alpha helix and beta sheet distribution.

## Discussion

Nosocomial infectious diseases cause the death of many people in intensive care units worldwide. Among the bacteria that cause intensive care unit (ICU) infection, *A. baumannii* has a devastating impact due to having multiple antibiotic resistance[16]. Arsenal of antibiotics that target *A. baumannii* failing by time, there is an urgent need to develop vaccines against this species to protect critically-ill ICU patients[17–19].

Recently, two main strategies were followed to identify vaccine candidates against *A. baumannii*. One of them was immunoproteomic approaches which use blood sera of *A. baumannii* infected patients or animal models[20]. The other one was a computational method based on *in silico* analysis and prediction of antigenic structure by using bacterial genome sequences[21,22]. In the presented study our strategy was to capture and identify immune-dominant antigenic molecule of *A. baumannii* via IgGs from human subjects’ sera. Our experimental strategy was relied on the hypothesis that immunoglobulins of infected persons can interact with immunologically dominant bacterial proteins. Subtractive analysis of captured proteins with patient and control immunoglobulins revealed immunologically remarkable molecules, which trigger *de facto* response of human immune system against the antibiotic-resistant *A. baumannii*.

Identification of immunogenic proteins using serum immunoglobulins applied successfully in the last. Recently, gel-based immunoblotting methods followed by MALDI-TOF analysis were used in identification of vaccine candidate proteins[20,22]. In gel-based methods, denaturation of proteins which opposed to *in vivo* scenario, brings a drawback in antibody-antigen interactions. These approaches were insufficient to determine bacterial antigens in real case scenarios; therefore, we performed immunoprecipitation which allows to pull-down the proteins in non-denaturing conditions as much as possible.

Recently published other studies have identified several proteins which are different than ours and a great majority of those have focused on the outer membrane and outer membrane vesicles. OmpA which has an important function in biofilm formation and pathogenicity of *A. baumannii*, was studied on murine model. In that study, diabetic mice were vaccinated with recombinant OmpA (rOmpA) and two weeks later, mice were infected via the clinical *A. baumannii* isolate. Vaccination was found to prolong survival in mice and significantly reduce the bacterial load in tissues, as vaccination caused high titers of anti-OmpA antibodies associated with survival in mice[19]. Ata and Bap are the other proteins that were applied in immunization against *A. baumannii*, yet the prevalence and solubility of produced recombinant protein were the major challenges for vaccination[18,23]. OmpK and Ompp1 were described as potential vaccine candidates by a reverse vaccinology study published. Chiang and colleagues **(2015)** found that these antigens were highly immunogenic, as a result of the very high production of IgG antibodies after vaccination mice with 2×3 μg of recombinant antigens and then, they confirmed 60% protection in the murine pneumoniae challenge model with the *A. baummannii* and porcine mucin. Another study on clinically isolated strains identified several proteins like CarO-like porin, AdeK, TonB, OmpH, and BamABDE that could be regarded as diagnostic markers[24] as well potential protective agents.

Evolutionarily, human accommodates a huge number of bacterial species, some of which are opportunistic pathogens. Classical vaccine candidate identification methodologies usually mislead or prone to capture conserved proteins *i*.*e*., OMPs. By using a subtractive approach, we identified 34 unique proteins via infected patient sera, in both membranous and secretome of *A. baumannii*. Majority of those proteins were not deeply analyzed with classical approaches or *in silico* and was somehow disregarded as potential vaccine candidate (*e*.*g*., B0V885 / ABAYE3068)[21,25,26]. Taking into consideration of structure, transmembrane properties, epitope regions and solubility of identified proteins, we further analyzed nine of them as potential recombinant vaccine candidate.

One of these proteins was B0VC68 (Glutamate/aspartate transport protein) is a member of the ABC superfamily, previously described as a periplasmic protein with an increased expression in tetracycline response condition[27] and in acid tolerance[28]. Identification of B0VC68 specifically in antibiotic resistant strains and its relation with tetracycline response makes this protein a valuable target, not only for vaccine development but also for small molecule targeting studies. B0V4F6 (AdeC) was another outer membrane protein, functions as efflux transmembrane transporter activity. It has been reported that AdeC has an important role in drug efflux and extrusion of compounds[29] and was known as important in maintenance of antibiotic resistance. Ni and collegues[30] suggested the AdeC as a vaccine candidate in a reverse vaccinology study. Contradictory to its function as efflux pump and structural predictions, our initial analysis with common bioinformatical tools (BOCTOPUS, TMHMM and PROSO II), AdeC was found to be as a soluble protein without any transmembrane region. Further structural prediction by AlphaFold2 confirmed that this AdeC is a transmembrane protein and which cannot be considered as a candidate for recombinant soluble protein. B0VE52 had also been listed as soluble proteins by common prediction tools and it has been suggested as a candidate for *A. baumannii* vaccine *in silico* approaches[21]. Our structural prediction via AlphaFold2 lead us to that the B0VE52 was another unfortunate candidate with a beta-barrel structure. Relying on the classical linear sequence-based prediction approaches have potential to mislead outcomes and state of art 3-D structural prediction tools should be taken into the consideration.

Another protein identified from the secretome, BamD (B0V8H0) is a part of the membrane protein assembly complex involved in the insertion of beta-barrel proteins into the outer membrane. Although any study on the immune potential of BamD was not found in literature, previously it has been reported that immunization with BamA, partner of BamD in establishing membrane assembly, increased the survival to 60% in the murine model[31]. A0A0R4J8Q3 (FklB) protein that function as peptidyl-prolyl cis-trans isomerase was identified by our experiments. This membrane annotated protein has a role in protein folding process according to gene ontology analysis. FklB was reported previously as and vaccine candidate by Chiang and collegues[32] and proved that the protein is highly immunogenic. Like FklB, SurA (B0VD00) which has role in outer membrane protein folding and assembly was captured and identified in our study. Functional similarities with FklB[33] and interactions with BamA[31] may indicate that this protein is a potential vaccine candidate. B0V9Z6 was a putative PQQ-dependent aldose sugar dehydrogenase and it has oxidoreductase function was identified from secretion of A. *baumannii* has significant B-cell epitope region and according to our structural model predicted epitopes are located on the accessible part of the protein which makes it a likely protein vaccine candidate. Up to knowledge, there was no record as to the immunogenicity and protective role of this protein in any model.

One of the identified proteins from the membraneous structure by immunoprecipitation was B0VAB5 (Putative bifunctional protein), a large protein consisting of 1071 amino acids that have lytic transglycosylase and hydrolase activity. According to gene ontology annotations, this protein has a role in the peptidoglycan metabolic process. Having LysM repeat regions implies that this protein is a cell wall binding protein and locates on the accessible surface of the bacteria[34]. Epitope predictions provide us with the presence of a considerable number of epitopic regions. Overlapped epitope prediction and 3D-structural analysis showed us that predicted epitopes were placed on the accessible parts of protein which is crucial for a vaccine candidate. Up to our knowledge, this protein has not been reported as a vaccine candidate for *A. baumannii*. Although there is lack of evidence in literature about this entity, our experimental approach allowed us identification of B0VAB5 as an immunogenic protein that directly captured via human IgGs.

Among the identified proteins, B0V885 caught our attention by having the number of epitopic regions and physically being a part of secretome. Interestingly, it was the only protein that can be immunoprecipitated directly from conditioned media without further enrichment. This suggest that B0V885 was highly abundant and immunodominant protein, therefore easy to detect via immunoproteomics. B0V885 (Putative membrane protein exposed to the bacterial surface) consisting of 974 amino acids. Epitope analysis of B0V885 presented that the protein has 8 significant mostly repeated epitope regions between 79^th^ and 656^th^ amino acids. Structural evaluations by AlphaFold2 were showed us this protein was consisting of mostly β-Sheets, and our predicted epitope regions fall within this structurally unique part rather than relatively small, helical part of the protein. When the whole amino acid sequence analyzed the solubility of intact protein was below solubility threshold according to PROSO II, nevertheless a partial analysis of repeated epitopic regions was found to be soluble. Furthermore, GlycoPP tool suggested that those repeated units have high tendency to O-Glycosylation, which might enhance the antigenicity and solubility of a protein. Presence of most likely an insoluble protein in the secretome might be evidence that this protein could be carried out via extracellular vesicles. This phenomenon might also explain why this protein only captured from naïve conditioned media. Interestingly, the comparison of the reference sequences of B0V885 between antibiotic-sensitive and resistant strains revealed the presence of an additional 17 amino acids residue in the resistant strain. Presence of this peptide sequence might affect the unknown function of this protein and it might be related to drug resistance. Further studies on this protein structure, function and its relation with drug resistance are needed.

In conclusion, after comprehensive proteomics and bioinformatic analyses, our study provided data about remarkable vaccine candidates. Capturing and identification of the bacterial proteins with patient immunoglobulins provides *de facto* vaccine candidates since these molecules can trigger immune response and could establish immunity against pathogen.

## Material and Methods

### Blood sample collection

Blood samples *of A. baumannii* bacteremia diagnosed patients, kept in the intensive care unit of the Erciyes University Hospital, were collected with the approval of the local Human Ethical committees (ERU_ LEC_2013-445), and with the written consent of the patients. Twenty-nine infection-positive cases, non-infected thirteen patients as the negative control of Intensive Care Unit (ICU) and three as the external control group from healthy individuals were used for this study. 10 mL of the whole blood were collected into Vacutainer sample tubes and centrifuged at 3000 rpm for 10 min. Serum samples were stored in -80 °C for further analysis.

### Bacterial strain, growth conditions, and protein sample collection

Multiple drug-resistant (BAA-1710) and non-resistant (ATCC 17978) *A. baumannii* strains were purchased from ATCC. Well-isolated colonies of both strains were grown in Luria-Bertani broth (LB) overnight at 37 °C on an orbital shaker 200 rpm per minute. 1:100 (v/v) dilution of the culture was started and grown until OD_600_ 1.0-1.2 were caught. Suspension of bacteria was chilled on ice and centrifuged at 8000 x *g* for 20 min at 4 °C. To perform immunoproteomic analysis supernatants were used for secretome analysis and the pellets were subject to outer-membrane protein (OMP) isolation.

### Naive Secretome protein isolation

Secreted protein was harvested as mentioned before and to increase the coverage of the analysis and obtain the secreted proteins in their natural confirmation, conditional media was directly used immunoprecipitation. MultiMACS protein A/G microbeads (Miltenyi Biotec, Germany) were used to catch the secreted proteins as described by[35].

### Phenol Extraction Method

Conditional media of *A. baumannii* were collected by centrifugation and 1:8 (v/v) phenol (Sigma Aldrich, Germany) was added on. After 30 minutes incubation at 4 °C, samples were centrifuged at 6000 x *g*. Protein containing phenol phase was transferred to new collection tube, then equal volume of Back Extraction Buffer (0.1 M Tris HCl pH:8, 20 mM KCl,10 mM EDTA, 0.4% β-mercaptoethanol) were added on samples[36]. After 15 minutes incubation at room temperature, samples were centrifuged at 6000 x *g* for 15 minutes. Previous step was repeated and phenol phase was transferred to new tube. Precipitation buffer (0,1 M NH_4_OAc in Methanol) was added to sample at 5:1 (v/v) ratio and samples were incubated at -20 °C overnight. the supernatant removed carefully, 1 mL of precipitation buffer was added on pellet and incubated -20 °C for 20 minutes. Following the centrifugation at 15000 x *g*, 4 °C for 30 minutes, pellet was washed twice with 80% acetone and dried. To perform immunoprecipitation samples resuspended with Cell Lysis Buffer (150 mM NaCl, 50 mM Tris-HCl pH:8, 0.1% Triton X-100)

### Outer-membrane protein isolation

Bacterial pellets were obtained by using centrifugation at 5000 *x g*, 4 °C for 10 min, washed twice with DPBS and stored -80 °C overnight. Frozen pellets were thawed and resuspended with buffer (10 mM Tris-HCl pH:7.5). Protease inhibitors were added on suspension and sonicated for 10 cycles (15 seconds sonication 45 seconds cooldown). The centrifugation process was repeated to get rid of the cytosolic contaminants. The ultracentrifugation process was performed according to a method described by[37]. Samples were centrifuged at 108000 x *g* and 4 °C for 15 minutes. 2% Triton X-100 containing Tris-HCl solution was added on pellet and incubated at room temparature. Following the incubation samples were centrifuged with the paramaters indicated in the previous step. Membrane proteins containing pellet were dissolved in cell lysis buffer (CLB) for immunoprecipitation.

### Immunoprecipitation

In order to increase the coverage of analysis beyond the limitations of classical immunoblotting methods, the immunoprecipitation approach which makes it possible to obtain the proteins in their naïve, confirmational forms were used. Collected secretome and isolated membrane proteins were incubated with patients’ sera and also control sera for one hour at 4 °C, then 50 μL of protein G conjugated nanobeads (Miltenyi Biotec, Germany) were added on samples. At the end of the 30 minutes, incubation samples were applied on paramagnetic columns that were washed with cell lysis buffer. Following the passing of samples, columns were washed four times with lysis buffer previously, then washed with a low-salting solution to get rid of the salt and detergent residues. Labeled antigenic molecules were torn off from columns with pre-warmed Laemmli Buffer that contains 50 mM DTT. Eluents were collected for further step of sample preparation.

### Filter Aided Sample Preparation (FASP) of immunoprecipitated proteins

FASP Protein Digestion Kit (Expedeon, UK) was used in the sample preparation process. Following the immunoprecipitation, 100 μL eluent was mixed with urea sample solution, transferred on spin filter and centrifuged at 14000 x *g* for 15 minutes. After this step, 200 μL urea sample solution was added on spin filter and centrifugation step was repeated with previous parameters. Alkylation of cysteine residues have been done by incubating with 100 μL iodoacetamide and samples were centrifuged at 14000 x *g* for 10 minutes. Alkylated samples were washed three times with urea sample solution and after the collection tube has been emptied additional three washes were performed with ammonium bicarbonate. 400 ng/mL Trypsin/Lys-C (Promega, USA) were added on spin filter and incubated overnight at 37 °C. Following the Trypsin/Lys-C digestion, 400 ng/mL Trypsin-Gold (Promega, USA) were applied on spin filter and samples incubated additional six hours at 37 °C. Subsequently to digestion process, spin filters placed on new collection tube. 50 mM ammonium bicarbonate was added on spin filters and samples were centrifuged. Then samples on spin filter were eluted by 500 mM NaCl solution by centrifugation at 14000 x *g* for 10 minutes. Collected samples were dried with vacuum concentrator (Speedvac, Eppendorf, USA).

### ZipTip Purification of Samples

In order to remove the salts from samples and purify them ZipTip purification method was used. In this step ZipTip C18 (Merck-Millipore, Germany) tips were used. Digested peptides were resolved with 10 μL 0,1% TFA (Sigma, Germany) containing mass spectroscopy grade water (Fluka, USA). Before sample purification, C18 materials of ZipTips were washed with 10 μL ACN, conditioned twice with 70% ACN and equilibrated twice by using 10 μL of 3% ACN, and 0,1% acetic acid. Afterward resolved samples were pipetted 10 times for each sample by using equilibrated ZipTip. Following this step, captured peptides in C18 material were washed twice by pipetting of 10 μL 5% ACN, 0,1% acetic acid and in this way, impurities were washed out. Purified peptides were eluted from ZipTip by using 10 μL 60% ACN, 0,1% acetic acid and dried by using vacuum concentrator. Prior to analysis samples were resolved in 10 μL of 3% ACN, 0,1% formic acid and transferred to thinglass placed in MS vials.

### Mass Spectrometry Analysis

Shotgun proteomic analyses have been performed with AB SCIEX TripleTOF® 5600+ integrated to LC-MS/MS Eksigent ekspert™ nanoLC 400 System (AB Sciex, USA). NanoLCcolumn, 3 μm, ChromXP C18CL and nanoAQUITY UPLC® column (1,8 μm HSS T3 75 μm x 150 mm) was used at trap-elute mode for the separation of peptides. Within the 310 minutes analysis time, Data Dependent Acquisition (DDA) Top20 tandem MS were performed and raw data analysis and multiple analytical data measurements in a single sample were done by using Analyst® TF v.1,6 (AB Sciex, USA) software. Precursor and production evaluations were completed with PeakView (1.2, AB Sciex). Generated peak-lists that contain MS and MS/MS spectra were used in identifying proteins, protein isoforms and their modifications with ProteinPilot 4.5 Beta (AB Sciex, USA). Identification was done by using the UniProtKB based reference library of resistant strain *A. baumannii* BAA 1710 (UP000002446) and non-resistant strain *A. baumannii* 17978 (UP000006737). During identification of proteins false discovery rate (FDR) was determined 1% and proteins that contain at least two identified peptide fragments were considered as correct identification.

### Bioinformatic Evaluations

Identified peptides were grouped with Venn diagrams. Linear B Cell Epitope Prediction tool of Immune Epitope Database (IEDB) was used to predict the epitope from identified amino acid sequences[38]. Bepipred Linear Epitope Prediction method was used, window size was determined as seven amino acids during calculations[39]. Beta-barrel structure of proteins was analyzed by Protein Homology/analogY Recognition Engine 2.0 (PHYRE2) web-based software[21]. BOCTOPUS database was used to determine both beta-barrel structure and transmembrane properties of proteins[40]. As another transmembrane site determining web-based tool TMHMM 2.0 (Transmembrane Hidden Markov Model) was also used to analyze the transmembrane sites of proteins [41]. In order to analyze the solubility of proteins PROSO II software was used[42]. Potential glycosylation of nine selected proteins were predicted by using GlycoPP v1.0 web server[43].

Fasta sequences of 8 intact proteins and one with the missing region (B0V885) were submitted to Alphafold v2.1.0[44] to generate the 3D structure modellings of the respective proteins with default parameters. We had selected three models for each run to get the best ranking models. Python script provided by Mayachemtools[45] was used to produce Ramachadran plots of the predicted structures. We performed all of the computations on our local HPC cluster.

## Acknowledgements

This work was supported by The Scientific and Technological Research Council of Turkey (TUBITAK) (Project Number: 114S571). We would like to thank Soriene Nazik OZCAN and Elif Abagail OZCAN for language editing.

## Author contributions

Study design: S.O., M.D., A.U.K., G.D., S.A.G., H.G., M.B.A. Blood sample collection: M.D., A.U.K., Sample Preparation: S.O., S.A.G., M.B.A. Proteomic Analysis: S.O., S.A.G., M.B.A., H.G. Bioinformatic evalutions S.O., S.A.G., M.B.A., H.G. Manuscript preparation: S.O., M.D., A.U.K., G.D., S.A.G., H.G., M.B.A.

## Figure Legends

**Table 1**. List of 34 identified resistant *A. baumannii* outer membrane and secretome proteins, their molecular mass (DA), amino acid lengths, encoding genes, localizations in the cell and annotated functions.

## Supplementary File Legends

**Supplementary Figure 1**. Immune Epitope DataBase (IEDB) results of nine selected proteins. Analysis was done by using default parameters of B-Cell epitope prediction tool Bepipred Linear Epitope Prediction 2.0. Threshold value was 0.35, window size was chosen as seven means center position four.

**Supplementary Figure 2**. Improved topology prediction of transmembrane beta-barrel structure of the candidate proteins were depicted by BOCTOPUS 2. Default parameters were used.

**Supplementary Figure 3**. TMHMM posterior probabilities results of nine selected proteins. Default parameters were used, outputs of analysis gathered extensive, with graphics.

**Supplemantary Table 1:** Solubility scores of identified 34 proteins were determined via PROSO II. Default parameters were used, solubility threshold value was set to be 0.6.

**Supplementary Table 2**. Potential O-Linked glycosylated sites of candidate proteins were predicted via GlycoPP. Predictions based on Binary Profile of Patterns and SVM (Support Vector Machine) threshold was used as default.

---

**Graphical Abstract.**
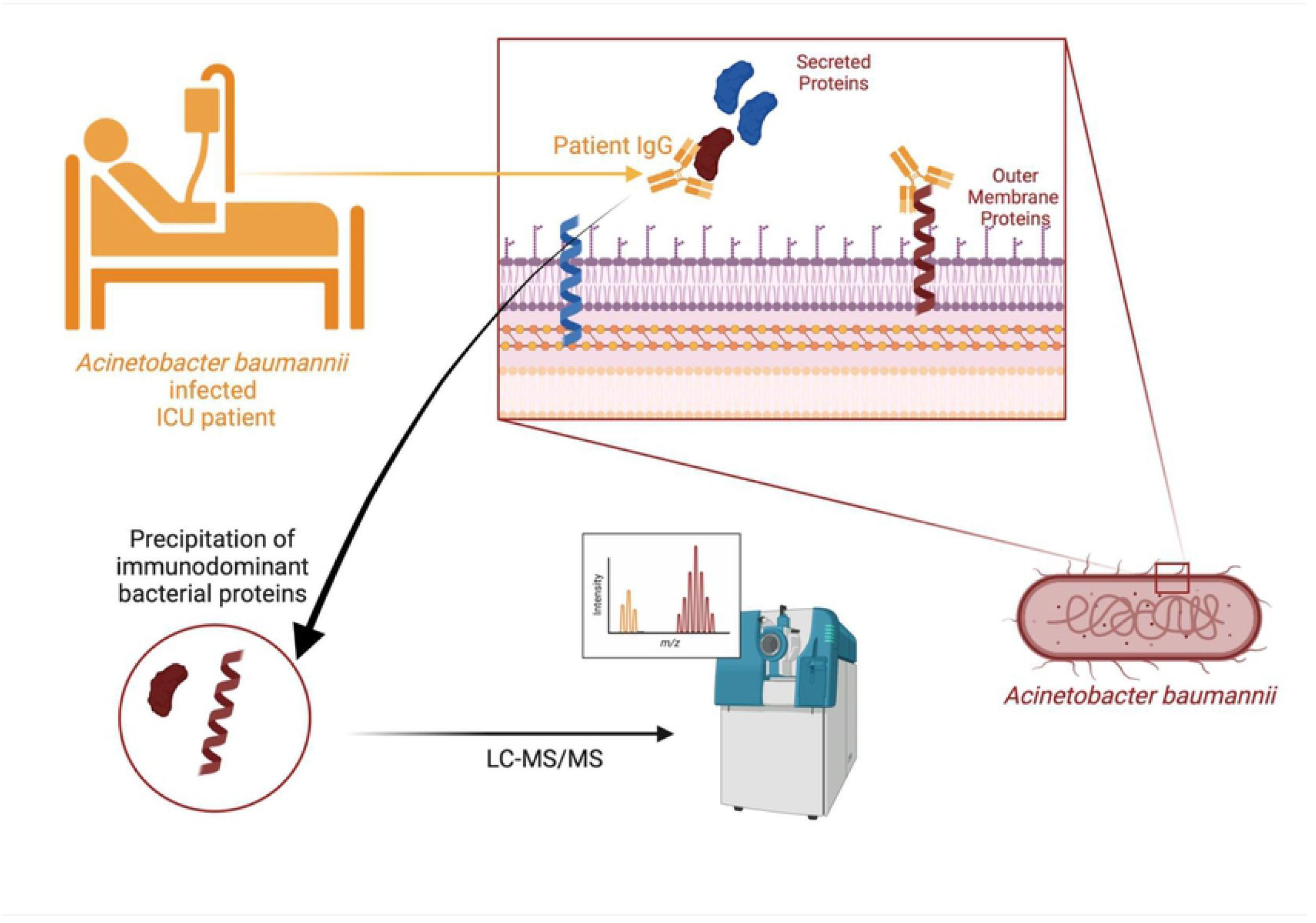

